# Laniakea: an open solution to provide Galaxy “on-demand” instances over heterogeneous cloud infrastructures

**DOI:** 10.1101/472464

**Authors:** Marco Antonio Tangaro, Giacinto Donvito, Marica Antonacci, Matteo Chiara, Pietro Mandreoli, Graziano Pesole, Federico Zambelli

**Affiliations:** Institute of Biomembranes, Bioenergetics and Molecular Biotechnologies, National Research Council (CNR), Via Amendola 165/A, 70126 Bari, Italy; National Institute for Nuclear Physics (INFN), Section of Bari, Via Orabona 4, 70126 Bari, Italy; Department of Biosciences, University of Milan, via Celoria 26, 20133 Milano, Italy; Department of Biosciences, Biotechnologies and Biopharmaceutics, University of Bari, Via Orabona 4, 70126 Bari, Italy

## Abstract

**Background:** Galaxy is rapidly becoming the de facto standard among workflow managers for bioinformatics. A rich feature set, its overall flexibility, and a thriving community of enthusiastic users are among the main factors contributing to the popularity of Galaxy and Galaxy based applications. One of the main advantages of Galaxy consists in providing access to sophisticated analysis pipelines, e.g., involving numerous steps and large data sets, even to users lacking computer proficiency, while at the same time improving reproducibility and facilitating teamwork and data sharing among researchers. Although several Galaxy public services are currently available, these resources are often overloaded with a large number of jobs and offer little or no customization options to end users. Moreover, there are scenarios where a private Galaxy instance still constitutes a more viable alternative, including, but not limited to, heavy workloads, data privacy concerns or particular needs of customization. In such cases, a cloud-based virtual Galaxy instance can represent a solution that overcomes the typical burdens of managing the local hardware and software infrastructure needed to run and maintain a production-grade Galaxy service.

**Results:** Here we present Laniakea, a robust and feature-rich software suite which can be deployed on any scientific or commercial Cloud infrastructure in order to provide a “Galaxy on demand” Platform as a Service (PaaS). Laying its foundations on the INDIGO-DataCloud middleware, which has been developed to accommodate the needs of a large number of scientific communities, Laniakea can be deployed and provisioned over multiple architectures by private or public e-infrastructures. The end user interacts with Laniakea through a front-end that allows a general setup of the Galaxy instance, then Laniakea takes charge of the deployment both of the virtual hardware and all the software components. At the end of the process the user has access to a private, production-grade, yet fully customizable, Galaxy virtual instance. Laniakea’s supports the deployment of plain or cluster backed Galaxy instances, shared reference data volumes, encrypted data volumes and rapid development of novel Galaxy flavours, that is Galaxy configurations tailored for specific tasks. As a proof of concept, we provide a demo Laniakea instance hosted at an ELIXIR-IT Cloud facility.

**Conclusions:** The migration of scientific computational services towards virtualization and e-infrastructures is one of the most visible trends of our times. Laniakea provides Cloud administrators with a ready-to-use software suite that enables them to offer Galaxy, a popular workflow manager for bioinformatics, as an on-demand PaaS to their users. We believe that Laniakea can concur in making the many advantages of using Galaxy more accessible to a broader user base by removing most of the burdens involved in running a private instance. Finally, Laniakea’s design is sufficiently general and modular that could be easily adapted to support different services and platforms beyond Galaxy.

## Background

The recent improvements in our capacity to gather vast amounts of complex, multi-layered and interconnected biomolecular data demand a parallel development and enhancement of the computational tools that we employ to analyse and handle this wealth of information. On the other hand, the rapid proliferation of those tools can render the execution of complex bioinformatics workflows cumbersome due to, among other things, incompatible data formats, long and convoluted command lines, versioning, and the need to handle and store a multitude of intermediate files. In turn, that not only makes harnessing the information contained in the biomolecular data unnecessarily onerous even for expert bioinformaticians but represents also a significant obstacle to reproducibility of the analyses [1] as well as an intimidating barrier for biologists aspiring to explore their own data in autonomy [2], students [3], and health-care providers adopting *clinical bioinformatics* approaches within their medical protocols [4]. For these reasons, in the past few years, a considerable effort has been put in the development of workflow manager platforms for bioinformatics (see [5] for a review). They usually provide integrated interfaces that not only provide a more user-friendly work environment but improve reproducibility, facilitate data sharing and enable collaborative data processing.

### Galaxy

The Galaxy platform is one of the most successful examples of such workflow management systems. Indeed, by providing a consistent, user-friendly, flexible, effective and customizable gateway to a vast array of bioinformatics software and analysis workflows, Galaxy has attracted a vast and thriving community of users [6]. The software consists of an open source server-side application accessible through a simple web interface that serves as a gateway to a wealth of tools for datasets handling and analysis, workflow design, visualization and sharing of results. Although from the user’s perspective using a Galaxy service to run bioinformatics analyses is pretty straightforward, the deployment of a production-grade Galaxy instance requires the configuration and maintenance of an extensive collection of helper components (e.g. database management system, web server, load balancer, etc…) and an even more extensive collection of bioinformatic tools and reference data. These issues, coupled with the need of an adequate or dedicated IT infrastructure, required to properly support the Galaxy service for any non-trivial amounts of users or workloads, has usually restricted the role of Galaxy service providers to institutions or groups with suitable IT facilities and the appropriate technical know-how. At the time of writing, more than 125 Public Galaxy instances are available (https://galaxyproject.org/use/), serving a vast community of users [6]. Despite being useful and popular, the public nature of those services implies some hardly addressable shortcomings like, for example, limited quotas for computing and storage resources, lack of customization options and potential concerns for data security and privacy, a worry particularly noticeable when processing sensitive data. These considerations can, in turn, limit or outright interdict the usage of Galaxy public instances for some specific applications or categories of users, e.g., analyses requiring big or huge computing workloads or precision medicine researchers and operators.

### Cloud solutions to Galaxy provision

The cloud computing model [7] is rapidly gaining popularity within the life sciences [8–11] and the biomedical [4,12,13] communities. Among other advantages, it offers a set of solutions and features that can overcome or mitigate the drawbacks of Galaxy public instances described above. At the time being, several efforts have already been put forward in this regard. Globus Genomics [14] provides a Galaxy-based bioinformatics workflow platform, built on Amazon cloud services, for large-scale next-generation sequencing analyses. CloudMan [15,16] allows individual researchers to deploy Galaxy instances relying on arbitrarily sized compute cluster on the Amazon cloud infrastructure (it can also support OpenStack and OpenNebula through custom deployments). The Genomic Virtual Laboratory (GVL) [17], offers Galaxy through a middleware available on Nectar, the Australian cloud infrastructure for research, and also on the Amazon cloud. PhenoMeNal [18] is a recent effort to develop a Cloud Research Environment that employs Galaxy to provide domain-specific tools for metabolomics. Finally, Krieger and colleagues describe a possible configuration stack to deploy Galaxy on an OpenStack based IaaS (Infrastructure as a Service) [19]. The appeal of Galaxy cloud solutions is also made evident by the 2016 Galaxy update [20] reporting that over 2,400 Galaxy servers were launched on the Amazon cloud in 2015 alone, pointing to strong demand for ready-to-use but private virtual Galaxy instances. The Amazon Galaxy service [21] is, however, a commercial solution that can discourage researchers and research or healthcare facilities from adopting it, due to funding or budget issues, ethical concerns or legal requirements (e.g., EU General Data Protection Regulation). At the same time, other solutions like access to GVL through the Nectar cloud are not usually within reach of European researchers and are not completely interoperable with other resources available to them through European e-infrastructures. In this article, we introduce Laniakea, a software framework for the provision of *on-demand* Galaxy instances. Laniakea is devised to run on existing scientific e-infrastructures by leveraging the open and modular middleware architecture developed within the INDIGO-DataCloud H2020 project [22,23] (https://www.indigo-datacloud.eu/) and is at the same time simple to deploy and maintain by infrastructure managers and convenient for end-users.

### Disambiguation of the terms “user”, “Galaxy user” and “IaaS administrator”

In this work, we use the words “user” or “end-user” to indicate anyone owning an account to access an instance of Laniakea. An authenticated Laniakea user can deploy Galaxy instances and has administrator rights both over the deployed instances and the virtual hardware hosting them. A single Galaxy instance can be accessed and used by many people, or “Galaxy users”, and there is no need for them to be also Laniakea users. Indeed, any Galaxy instance created by Laniakea receives a public IP and manages accounts in the same way of any other Galaxy instance, also supporting anonymous login if the administrator so decides. Finally, we use the terms “IaaS administrator” or “IaaS manager” for the person or entity running a cloud infrastructure that offers Laniakea as one of their services.

## Methods

### INDIGO-DataCloud middleware components

Laniakea is based on the ElectricIndigo software release of the INDIGO-DataCloud H2020 project (INDIGO from now on), that officially ended in September 2017 and passed on its legacy to the EOSC-Hub, (https://www.eosc-hub.eu/), DEEP-HybridDataCloud (https://deep-hybrid-datacloud.eu/) and XDC (http://www.extreme-datacloud.eu/) projects. INDIGO aimed at making cloud e-infrastructures more accessible by scientific communities. In fact, the heterogeneity of the currently available scientific cloud infrastructures often entails portability issues between the different technologies employed that can result in the lack of one or more key features, e.g., automatic elasticity or multi-clouds deployments. In turn, these resources usually result in less efficient solutions than commercial counterparts relying on more homogeneous environments. INDIGO’s software catalogue [22] tackles e-infrastructures heterogeneity through comprehensive support to open standards and solutions, e.g., adopting both OpenStack (https://www.openstack.org) and OpenNebula (https://opennebula.org) as Cloud Management Platforms (CMPs) and allowing to transparently deploy applications using either virtual machines (VMs) or Docker containers. This result has been achieved by integrating the needs of a wide range of use cases and by taking advantage of publicly available open-source cloud components, adapting or enhancing them if needed to obtain the desired functionalities and embarking in the development of new software packages when strictly necessary. Fig. 1 provides a complete overview of the INDIGO PaaS software components and the installation sequence needed to deploy them correctly. Table S1 (in Supplementary2.docx) collects the URLs of all the INDIGO components required by Laniakea.

**Fig. 1.**
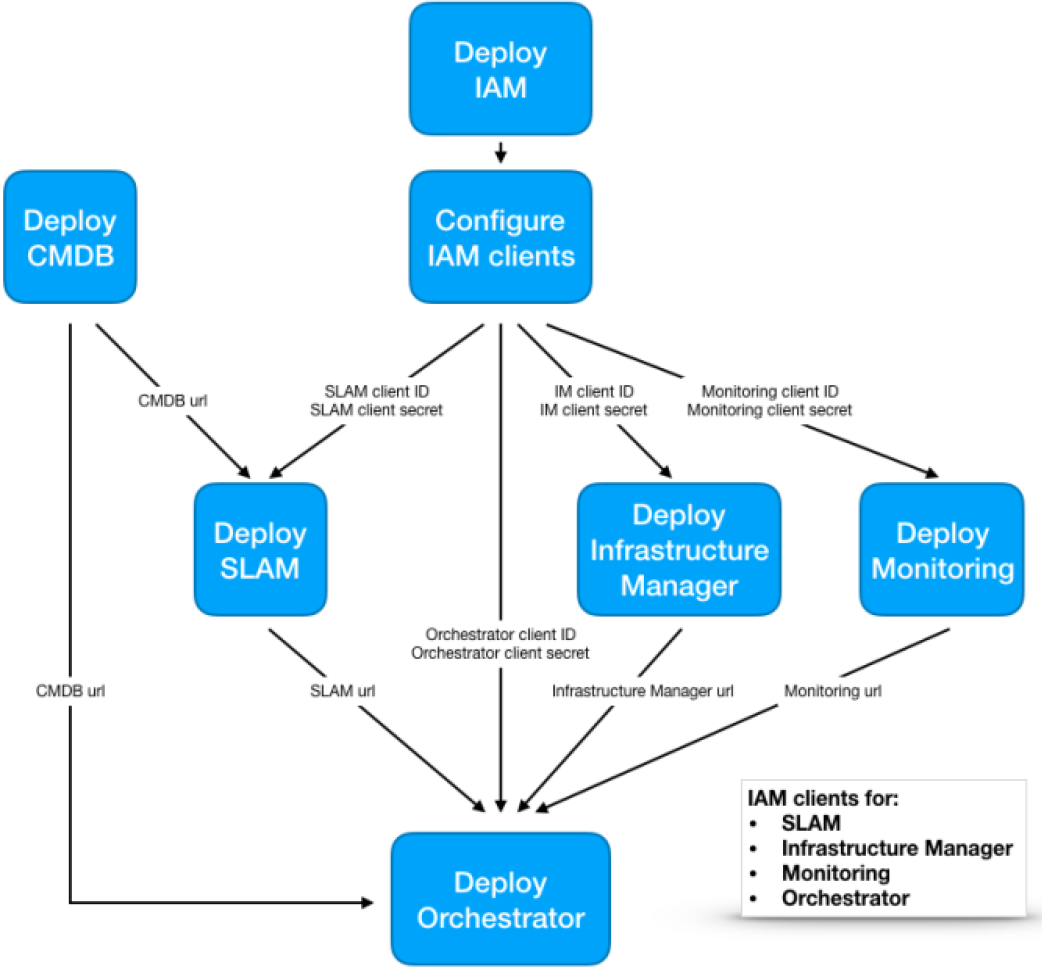
INDIGO-DataCloud PaaS software components and deployment procedure. The first step consists in installing and configuring IAM; this allows registered clients to request and receive information about authenticated end-users. Each INDIGO service must authenticate to a dedicated IAM client using a public identifier, the “client id”, and a private password, the “client secret”, provided during their configuration. In this way, IAM can verify that the connecting service is trusted and grants it the rights to interact with the other PaaS and IaaS services. The CMDB service installation follows as the second step; it provides detailed information on the available cloud sites and the virtual environments that they support. The following steps involve the deployment of SLAM, the Service Level Agreement Manager, the IM, and the Monitoring stack. Finally, the INDIGO PaaS-Orchestrator is deployed.

### Orchestrator and Infrastructure Manager

The INDIGO PaaS Orchestrator and the Infrastructure Manager (IM) components (https://github.com/indigo-dc/orchestrator), which are both based on the OASIS Topology and Specification for Cloud Applications (TOSCA) [24,25] open standard language, are in charge of setting up the virtual infrastructure environment and deploy the software framework. Cloud applications and services, together with their requirements and dependencies and regardless of the underlying platform, are described using YAML syntax [26]. This approach enables the deployment of complex applications from small reusable building blocks, the “node types”, and the topology of their relationships. The INDIGO PaaS Orchestrator coordinates deployment of the application according to the specifications described in the TOSCA template: it selects the most suitable cloud infrastructure site among those available and delegates the deployment and configuration of the virtual infrastructure on the target site to the Infrastructure Manager (IM) (https://github.com/indigo-dc/im). IM supports deployment on OpenStack, OpenNebula and commercial cloud providers alike, and serves as an abstraction layer for the definition and provision of the required resources. Finally, the software layout of the infrastructure’s nodes is described using Ansible roles (http://docs.ansible.com). They instruct the automation engine on how to install and configure the end-user applications or services, like Galaxy, on bare OS images. All in all, this architecture allows for the deployment of a wide range of cloud agnostic applications and services.

### Authorization and Authentication Infrastructure

The INDIGO Identity and Access Management (IAM) is an Authentication and Authorisation Infrastructure (AAI) service that manages users identities, attributes (e.g., affiliation and groups membership) and authorization policies for the access to INDIGO based resources. IAM supports several token-based protocols, i.e. X.509 [27], OpenID Connect [28] and SAML [29], thus allowing the federation of different infrastructures. This system allows users to connect to federated PaaS and manage heterogeneous and distributed resources through a single AAI service and with just one user account, easing the overhead experienced by end-users in case of multiple INDIGO services running on the same or federate infrastructures.

### Other INDIGO components

- The **CLUES** service is an elasticity manager for HPC clusters that enables dynamic cluster resources scaling, deploying and powering-on new working nodes depending on the workload of the cluster and powering-off and deleting them when no longer required. Once new jobs are submitted to the cluster Resource Manager (RM) queue (e.g., SLURM [30] or Torque [31]), CLUES contacts the Orchestrator and starts the provisioning of available temporary resources, thus improving the overall efficiency of the infrastructure.
- **FutureGateway** is built on top of the Liferay open source portal framework (https://www.liferay.com) and provides GUI based portlets that interact with the PaaS layer, allowing the authentication through the integrated INDIGO IAM portlets and the customization of relevant parameters of the TOSCA templates by the user before dispatching them to the Orchestrator for deployment.

### Encryption at block device level

The encryption layer is based on LUKS (Linux Unified Key Setup) [32], the current standard for encryption on Linux platforms. It provides robustness against low-entropy passphrase attacks using salting and iterated PBKDF2 passphrase hashing. LUKS supports secure management for multiple user passwords, allowing to add, change and revoke passwords without re-encryption of the whole device. Supplementary1.docx contains a detailed description of the Laniakea encryption framework.

### CernVM File System

CernVM File System (CVMFS) [33] is used to support common reference data repositories. CVMFS is a read-only POSIX file system originally developed to facilitate the distribution of High Energy Physics analysis software through HTTP protocol. Data files are hosted on any server and can be mounted concurrently on multiple compute nodes through a Linux filesystem module-based client (FUSE) that loads and caches only the specific portions of the files that are required, on-demand. This solution is suitable for quasi-static files that have to be shared across several geographically distributed clusters, and it supports local caches to speed up read operations on the hosted data too.

## Results

A wide range of Galaxy users and instance types do exist. For example, users may stretch from plain account owners to instance administrators and from developers to training courses teachers. At the same time, Galaxy instances can have different footprints in terms of computational resources, depending for example on typical usage or number of concurrent users; can be short-lived to satisfy temporary needs (e.g., a training course) or long-lived to provide a persistent work environment; public or private; needing advanced data security features in order to work on sensitive human data or not, etc…. As a consequence, Galaxy PaaS providers should be able to cover the requirements of the largest possible number of user/instance type combinations in order to maximize the convenience of their service. Laniakea’s architecture (Fig 2) has been developed to address all these heterogeneous needs swiftly, empowering end-users with a wide array of customization options that can deliver from out-of-the-box, stable, production-grade Galaxy instances already configured with collections of bioinformatics tools, reference data and cluster support, to blank-sheet Galaxy instances, ready to be completely personalized. Another key feature of Laniakea consists in the integration, for the first time at best of our knowledge, of a built-in technology to encrypt storage volumes on a Galaxy on-demand platform. This functionality can be used to provide robust data protection through state-of-the-art encryption protocols: the secure layer insulates stored sensitive data from unauthorized access both by malicious attackers or trusted users of the same cloud infrastructure and notably including the administrator(s) of the cloud and hardware layers themselves. Finally, Laniakea supports reference data sharing via CVFMS and a high degree of instance scalability through an array of deployment configurations ranging from single node Galaxy instances to SLURM managed Galaxy clusters, including also cluster elasticity based on CLUES.

**Fig. 2.**
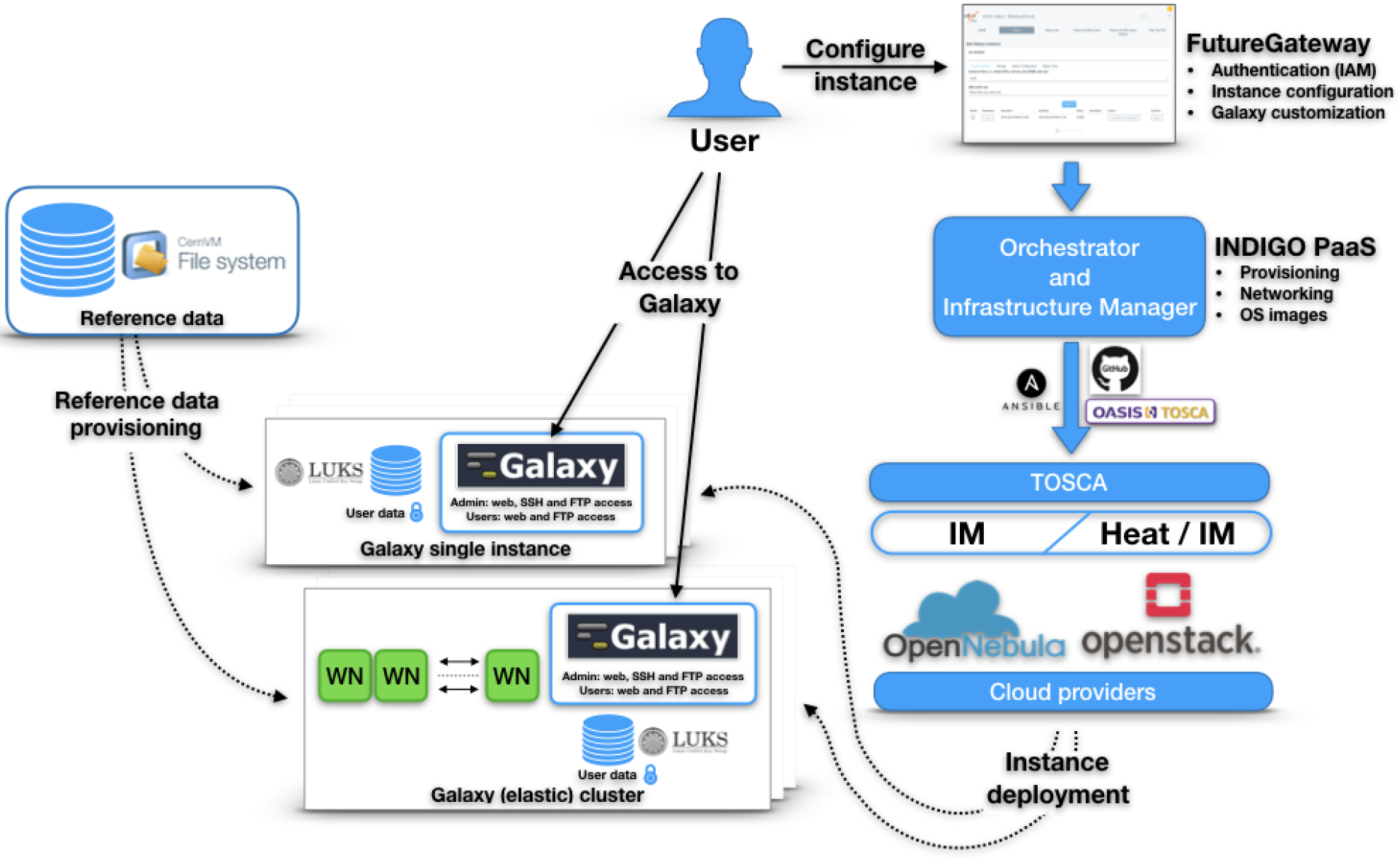
Architecture and deployment flow. The FutureGateway portal provides users with the front-end to configure and manage Galaxy instances, the resulting TOSCA templates are sent to the PaaS layer that employs INDIGO services to deploy instances over the IaaS, that in turn provides virtual hardware, storage, and networking. Finally, the Galaxy instance (single or cluster backed) is configured with the requested flavour and attached to a plain or LUKS encrypted storage volume (depending on user’s choice) and to the CernVM-FS shared volume with reference data. At the end of the process, the public IP address of the new Galaxy instance is provided to the user.

Table 1 provides a detailed features comparison table of Laniakea, PhenoMeNal and GVL. Table S2 (in Supplementary2.docx) collects the URLs of the software components of Laniakea. A video demo showcasing a typical user interaction routine is available at https://goo.gl/xnWNQd. In the following paragraphs, we describe in more detail the components and features of the software suite.

**Table 1.**
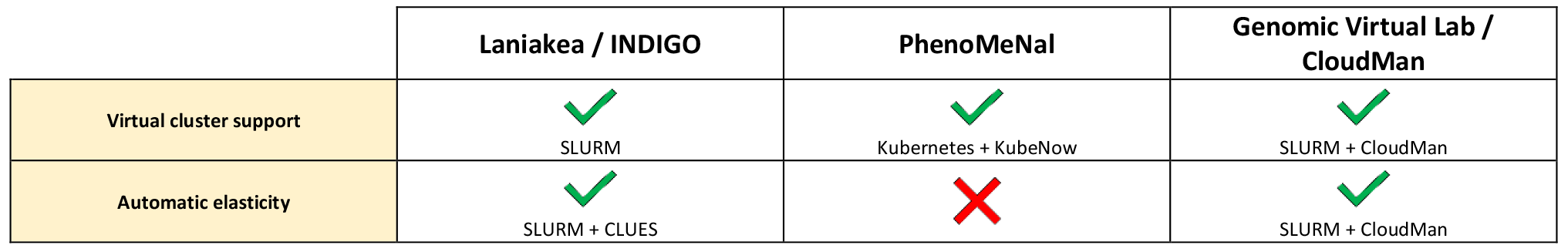
Comparative features table of Laniakea, PhenoMeNal and GVL. Virtual environment (light yellow background), Galaxy (light green background) and IaaS specific (light blues background) features are compared. Data for PhenoMeNal and GVL have been gathered from available documentation using our best effort. Only features related to the Galaxy on-demand platform have been considered, leaving aside GVL’s and Phenomenal’s support for other software (e.g., RStudio, Jupyter).

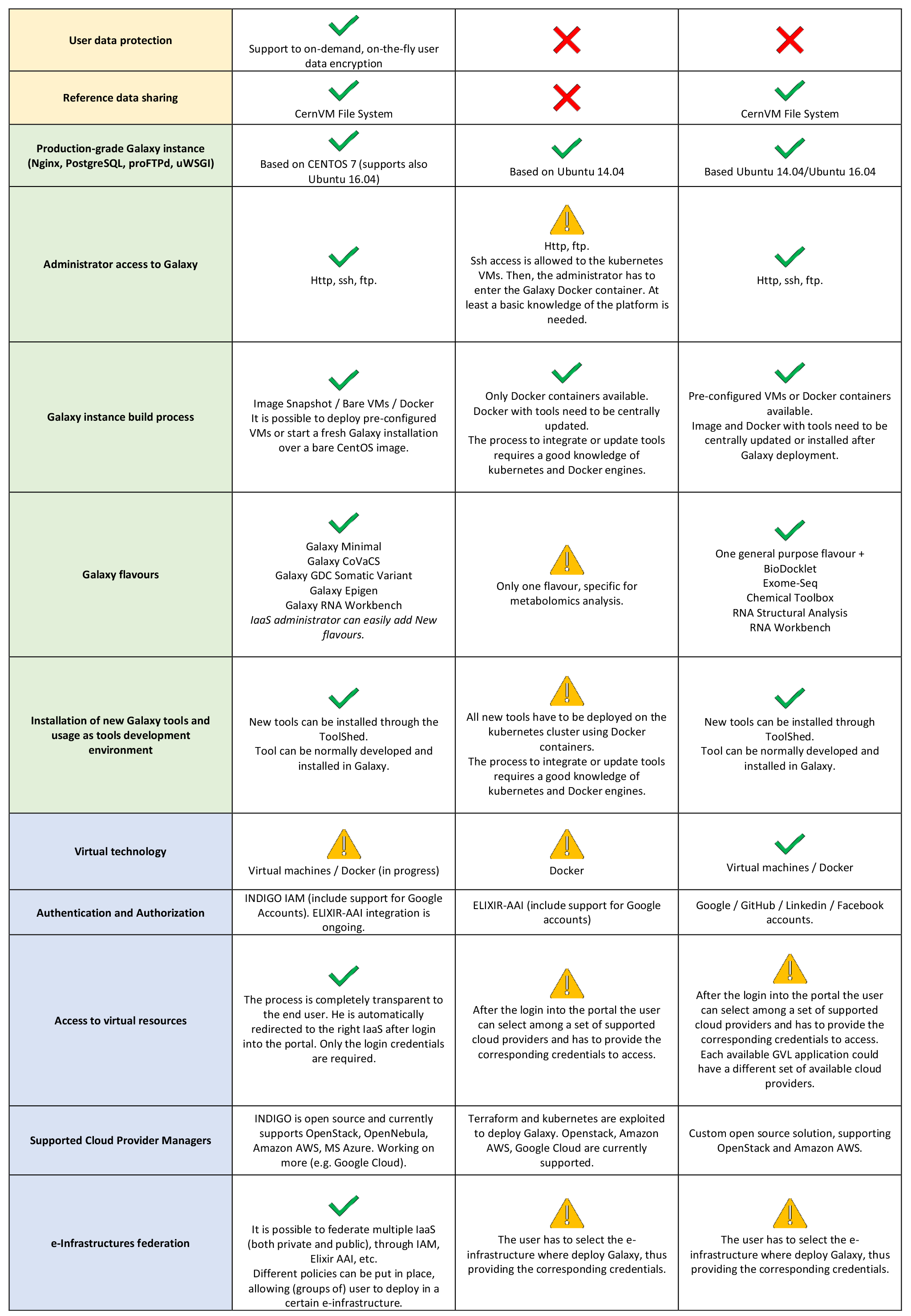

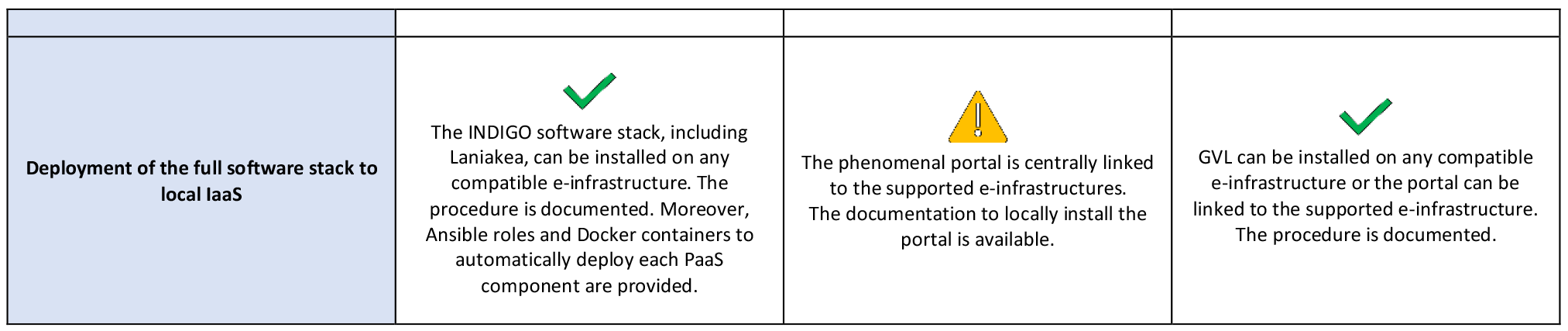

### The web front-end

The web front-end of Laniakea (Fig. 3), is built upon the INDIGO FutureGateway component and becomes accessible after authentication through INDIGO IAM. The front-end represents the entry point to the PaaS from a user’s point of view. It provides the configuration interface for initialization and deployment of all the virtual hardware/Galaxy instance type combinations available through separate configuration panels for Galaxy single VM or cluster deployments.

**Fig 3.**
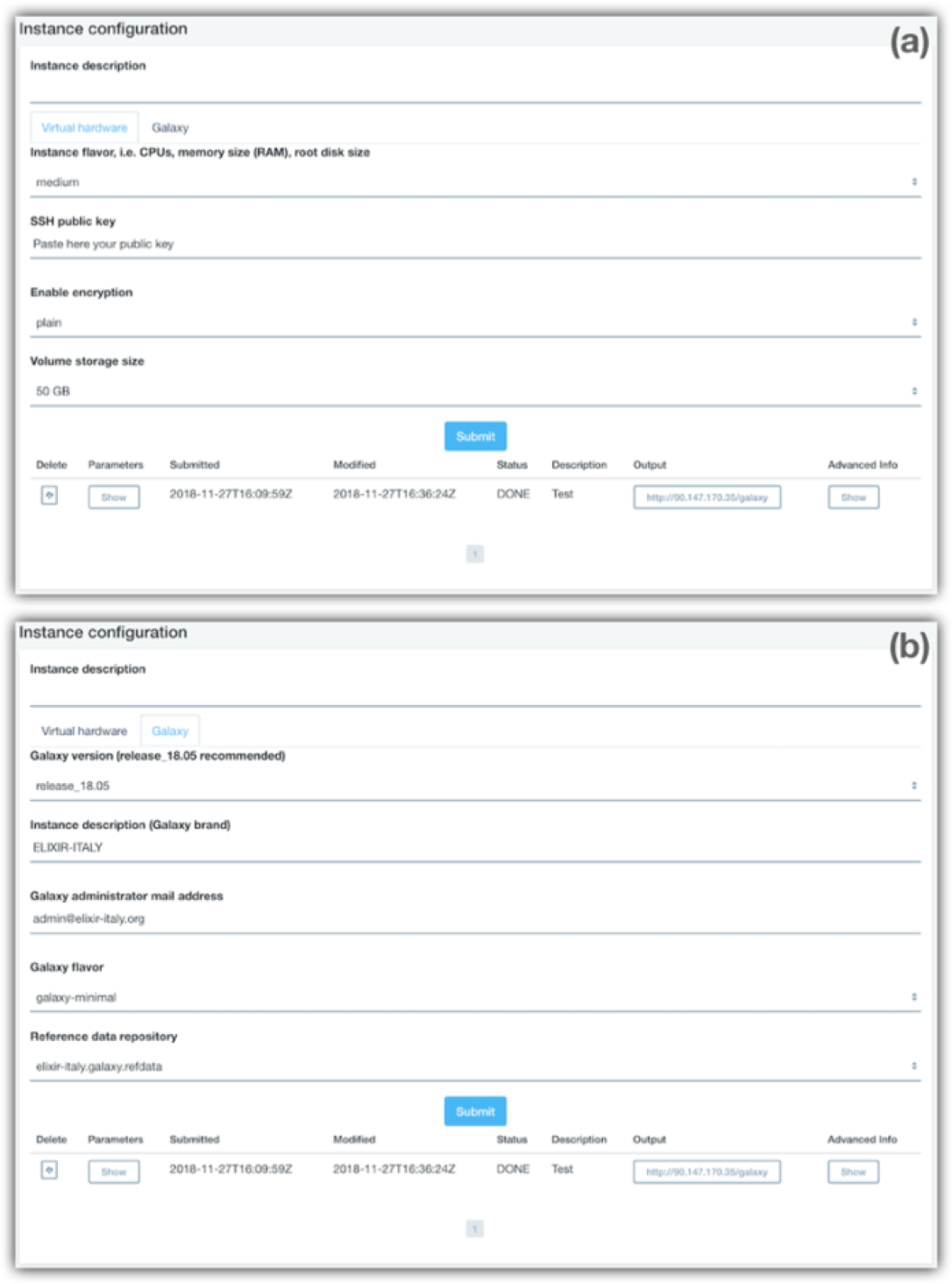
Web front-end. The web interface is organised using different tabs for each configuration task. The “Virtual hardware” tab (a) allows the selection of the virtual hardware, e.g., the number of CPUs and quantity of memory, storage size and to submit the user SSH public key that will grant access to the VM at the end of the deployment. For cluster deployments, the same tab also allows configuring the number of worker nodes required and their hardware configuration. The “Galaxy” tab (b) provides the software configuration options: Galaxy version, instance description, the mail address of the administrator, the flavour and Reference data repository from the available ones. The list of active instances and their status is displayed at the bottom of the page.

The configuration front-end is organized in two panels to guide users through the design of the new Galaxy instance:

- The Virtual Hardware configuration tab exposes the array of available hardware setups, differing for the number of virtual CPUs and quantity of RAM. Users are required to provide a valid SSH public key at this point. The key will be used to grant access to the Virtual Machine, once deployed. The storage volume size and type, i.e., plain or encrypted can also be selected at this stage. The Virtual Cluster deployment panel allows for the setup of both the front-end and worker nodes.
- The Galaxy Configuration tab allows the selection of the Galaxy release version among the ones currently supported, the reference dataset, the e-mail of the instance administrator and finally the *Galaxy flavour*. The Galaxy flavour is the pre-configured set of bioinformatics tools that will be installed and thus ready to use straight away after deployment (more on this in the “Galaxy flavours” section).

The settings defined through the interface are in turn submitted to the INDIGO Orchestrator that manages the deployment and configuration processes. When the procedure terminates, a public IP address becomes available which provides access to the freshly minted Galaxy server. From that moment onwards, the user has full administrator privileges over the instance and can access the underlying Virtual Machine through SSH using the private key corresponding to the public one that was provided. Public key authentication has been preferred over using passwords since this approach avoids most of the drawbacks of using and safeguarding passwords both for users and the IaaS administrator.

### The Galaxy environment

The deployment of a Galaxy multi-user production environment is a complex task that requires several auxiliary software components that in turn must be properly installed and configured. Laniakea automates this lengthy and error-prone procedure by spawning on-the-fly virtual environments. Their standard configuration (Table 2) is as follows: CentOS7 as the operating system (OS), PostgreSQL as the database engine, Nginx, uWSGI, and ProFTPD as the web, application and FTP servers respectively. Apart from the OS, for which there are no official recommendations, this configuration is rooted in the guidelines for production environments issued by the Galaxy Project itself (https://docs.galaxyproject.org/en/latest/admin/production.html). CentOS 7 has been selected as the reference OS for Laniakea virtual environments due to its well-known adherence to standards and the long span of the foreseen official support for this release (updates until June 30, 2024); however, Ubuntu 16 is also supported. Laniakea fully supports virtual machines while docker containers are currently in the final stage of testing (containers available at https://hub.docker.com/u/laniakeacloud/) and will be soon available, leaving the choice of the virtual environment solution to the preferences of the IaaS administrator. The on-the-fly deployment of the virtual environment offers a higher degree of agnosticism over pre-configured virtual machines and at the same time ensures that each new instance of Galaxy makes use of the latest available release of software components. However, this approach suffers from two possible limitations: first, the installation procedure takes time, about five hours on our test IaaS and second, there is a chance of some software components (mainly bioinformatics tools) failing to install due to any temporary unavailability of the online repositories needed for their installation. To overcome these limitations, Laniakea also provides a backup procedure to instantiate pre-configured images of Galaxy instances that the IaaS administrator can easily create from the on-the-fly ones previously described and provide to the user through the “Galaxy Express” configuration panel of Laniakea. In this latter case, the procedure will automatically adapt the configuration of the image to make it compatible with the virtual hardware and any other parameter (e.g., the instance administrator credentials) selected by the user. Apart from being more resistant to third parties repositories unavailability, this procedure allows faster deployments, i.e., less than an hour on our test IaaS, at the cost of a lesser degree of IaaS agnosticism.

**Table 2.** Laniakea’s Galaxy production environment components summary.

| Software component | Version installed by Laniakea |
| --- | --- |
| Operative System | CentOS 7 (supported Ubuntu 16.04) |
| Galaxy | release_17.05 and release_18.05 |
| PostgreSQL | 9.6 |
| NGINX | 1.12.2 |
| uWSGI | 2.0.17.1 |
| PROFTPD | 1.3.5e |

### Galaxy flavours

The Galaxy Tool Shed (https://toolshed.g2.bx.psu.edu/) is the official Galaxy tools repository and provides a convenient system for the administration and customization of a Galaxy web-server with tools and workflows tailored for its expected typical use. However, despite the significant improvements brought by the recent adoption of Conda (https://conda.io) for the management of packages and dependencies, the concurrent installation and configuration of many tools and their respective dependencies and reference datasets can still be burdensome and sometimes it requires a low-level understanding of the Galaxy environment and of the tools to be installed. For this reason, Laniakea provides a handy set of domain-specific *Galaxy flavours*, that is Galaxy instance enriched with cured collections of tools already installed, configured, tested, organised in workflows and ready to be used out-of-the-box. To access a target Galaxy instance and install the required bioinformatics tools Laniakea employs the official Galaxy Project library Ephemeris (https://github.com/galaxyproject/ephemeris), and YAML recipes are used to list the required packages. A helper Ansible role is used for the fine-tuning of packages configuration and to solve dependencies that, for any reason, are missing or malfunctioning upon installation with Conda. All in all, this approach allows IaaS administrators to easily prepare additional flavours by just creating new YAML recipes and eventually tweak the resulting configuration using the helper Laniakea Ansible role.

Currently, Laniakea provides five proof-of-concept flavours that are named “galaxy-minimal”, “galaxy-epigen”, “rna-workbench”, “GDC_Somatic_Variant” and “CoVaCS”. The first flavour is a bare instance providing just the default tools embedded in any Galaxy installation and provides a clear foundation for users who need a blank sheet ready to be customized to their own needs or for developers of new tools and flavours. The “galaxy-epigen” flavour is based upon the layout of the Italian Epigen Project’s (http://www.epigen.it/) Galaxy server (http://www.beaconlab.it/epigalaxy) and provides a selection of tools for the analysis of ChIP-Seq and RNA-Seq data. The “rna-workbench” flavour is based on [34] and includes more than 50 tools dedicated to RNA-centric analyses including, i.e., alignment, annotation, secondary structure profiling, target prediction, etc... The “GDC_Somatic_Variant” flavour is a porting of the Genomic Data Commons (GDC) pipeline for the identification of somatic variants on whole exome/genome sequencing data (https://gdc.cancer.gov/node/246), the resulting Galaxy workflow is shown in Fig. 4. In this case, Galaxy wrapper s for several tools, which were not previously available, have been developed from scratch. Finally, the CoVaCS flavour implements the homonymous workflow (Fig. S1 in Supplementary2.docx), described in [35], and set of tools for genotyping and variant annotation of whole genome/exome and target-gene sequencing data. All in all, we believe that these examples provide an adequate proof-of-concept of the relative easiness of generating Galaxy flavours for Laniakea.

**Fig. 4.**
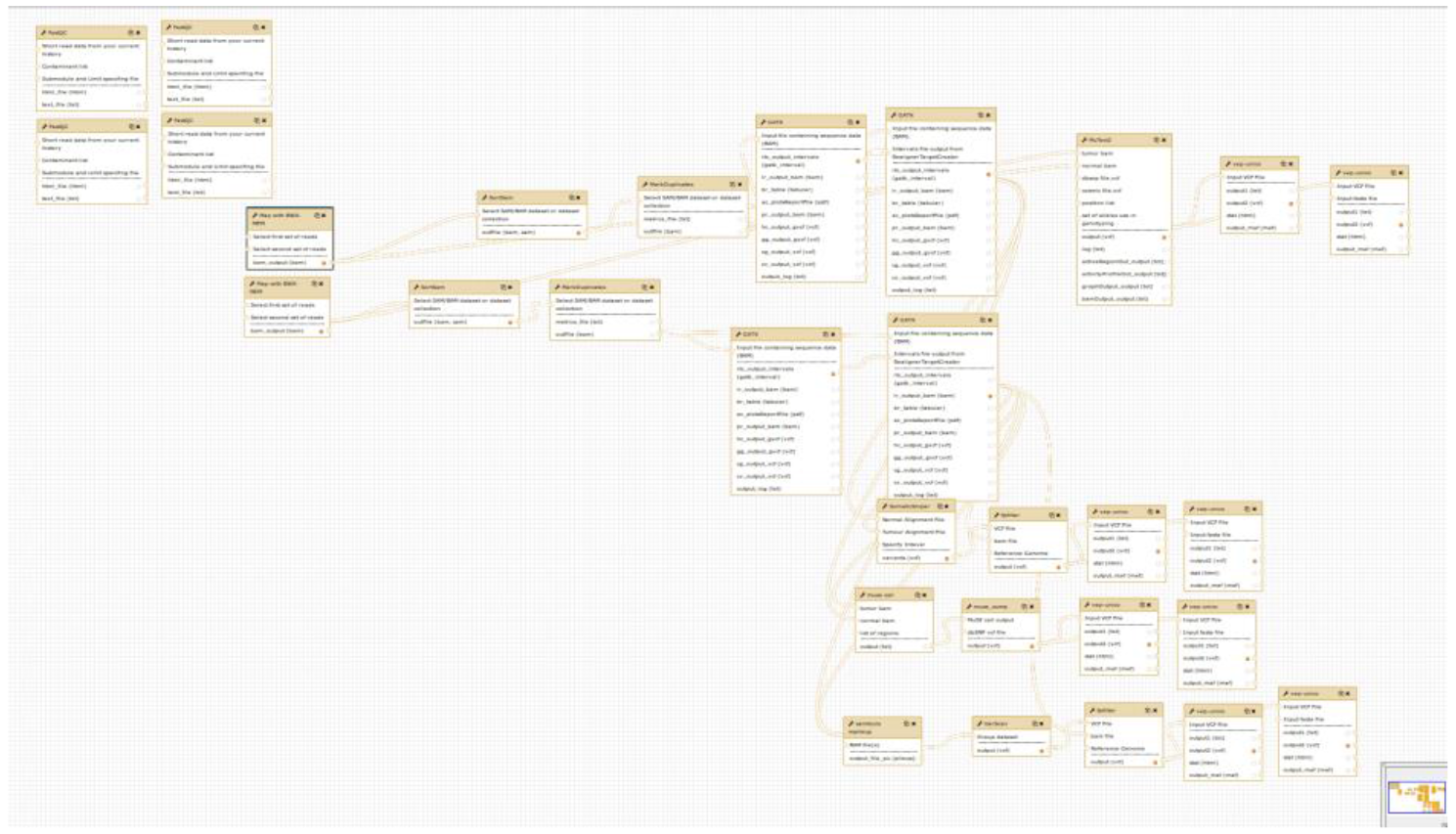
The GDC Somatic Variant analysis pipeline implemented as a Galaxy workflow for the corresponding flavour. The workflow design interface of Galaxy is a powerful instrument to elaborate complex workflows that link together the output and the input of different tools in an intuitive fashion.

### IaaS wide Reference data availability

To avoid useless replication of reference data and facilitate reproducibility of analyses, Laniakea employs a read-only CVMFS volume shared among all the instances of the same IaaS. The IaaS manager can choose to provide its own CVMFS customized reference dataset or to mirror or link a remote one, e.g., the one made available by the Galaxy Project. In the latter case, the required reference data need to be transferred from a remote service on-the-fly. To streamline the creation of new CVMFS repositories by the IaaS manager, we have made available a suitable Ansible role and documented the procedure. As a proof of concept, we provide three different repositories. The first is a manually curated reference dataset maintained by us, which contains the latest releases of the human, mouse, yeast, fruit fly and *A. thaliana* genomes and their corresponding indexes for Bowtie [36], Bowtie2 [37] and BWA [38,39]. The second is a repository tailored for variant calling in human, to be used with the CoVaCS and GDC Galaxy flavours, and provides access to a collection of publicly available variants, provided as VCF format files, derived from a selection of large-scale human genome resequencing projects [40–42], along with curated human genome assemblies and indexes obtained from the GATK bundle repository [43]. Finally, we provide a mirror of the Galaxy project “by hand” reference repository. In principle, this approach could be extended to optimize the use of storage resources and performances in case of a single user or group using several geographically distributed IaaS resources. Galaxy administrators needing additional reference data not included in these repositories will still be able to add them using the storage resources assigned to their instance.

### Data protection and isolation

Unless proper countermeasures are in place, data stored on a VM could be exposed to anyone with legitimate or illegitimate access to the underlying IaaS and physical hardware [44]. This issue poses technical, ethical and legal concerns, and in particular when sensible human genetic data are involved. These problems are exceptionally relevant for health operators and researchers involved in clinical bioinformatics or similar scenarios. We tackle this issue by providing to Laniakea users a security layer that seamlessly encrypts the storage volume using file system level encryption based on the *dm-crypt* Linux kernel’s subsystem coupled with LUKS encryption strategy based on aes-xts-plain64 cipher (see Table S3 in Supplementary2.docx for the detailed configuration parameters). The storage volume is encrypted using a *key stretching* approach: a randomly generated master key is encrypted using the user passphrase through PBKDF2 key derivation. This procedure makes both brute force and *rainbow tables* [45] based attacks more computationally expensive, and at the same time allows for multiple passphrases and passphrase change or revocation without re-encryption. Finally, the LUKS *anti-forensic splitter* feature protects data against recovery after volume deletion. The resulting instance layout consists of Galaxy running on top of a standard file system but transparently using the encrypted volume for storing data as long as it is unlocked and mounted (Figure 5).

**Fig. 5.**
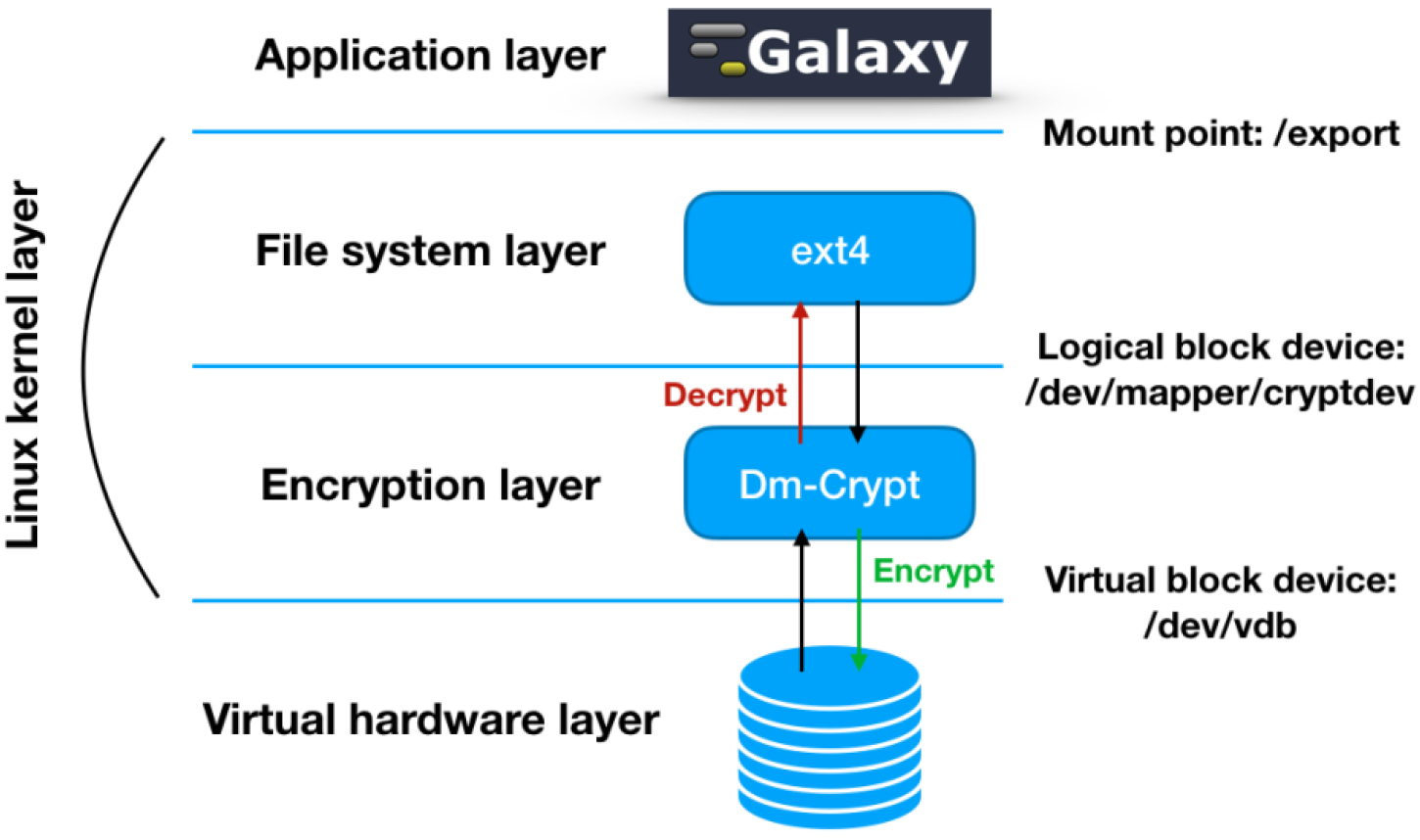
The relationship between Galaxy, the file system, and dm-crypt. Data are encrypted and decrypted on-the-fly when writing and reading through dm-crypt. The underlying disk encryption layer is completely transparent to Galaxy that employs a specific mount point in order to store and retrieve files from the volume.

The encryption procedure is coordinated by the IM which installs the encryption package and sends to the user an email containing the information needed to log in to the newly created VM, together with a brief description of the encryption procedure and a detailed step by step how-to (Figure 6). Users are thus required by this procedure to insert the passphrase for file system encryption manually through an SSH connection to the Galaxy instance being deployed; a similar system will allow mounting the encrypted volume each time the encrypted instance is re-booted. This two-steps solution has been devised to separate the orchestration of services from the encryption procedure, ensuring that the encryption passphrase is never being exchanged as plain text during the deployment procedure and avoiding any interaction with the IaaS administrator(s). As a result, the user of Laniakea is the only one holding the passphrase and thus able to unlock the encrypted volume.

**Fig. 6.**
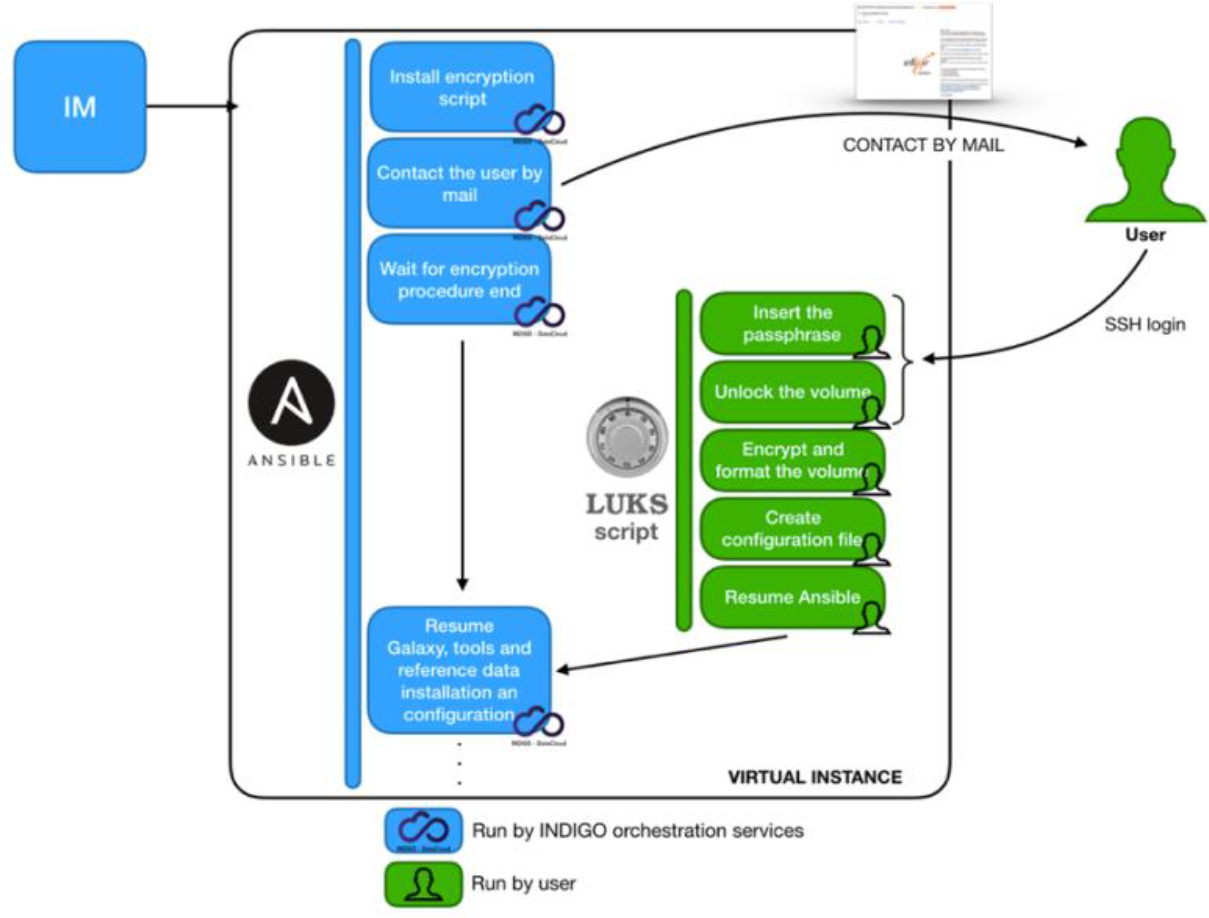
Storage encryption workflow. The user receives an e-mail with instructions to connect through SSH to the virtual Galaxy instance being created. The user is then requested to enter a passphrase that will be used to encrypt the volume and requested each time the encrypted volume will need to be unlocked, e.g. during a reboot of the VM hosting the Galaxy instance.

To validate our data encryption strategy, we simulated two different attack scenarios. In the first scenario, the attacker obtains unauthorized access to the unmounted encrypted volume, while the second simulates the improper use of administrator IaaS privileges when the LUKS volume is already unlocked and in use by a running Galaxy instance.

For the first scenario we compared two identical volumes, one encrypted and the other not, both attached to the same Galaxy instance, with the same set of permissions and each containing a copy of the same plain text file. Once detached, we created a binary image file of each volume and tried to access the data structure through *hex dump*. We were able to quickly retrieve the original content of the text file from the non-encrypted volume while the hex dump of the encrypted volume did not contain the original text in any discernible form.

For the second scenario, we tried to read data from the volume already mounted on a running Galaxy instance, using the OpenStack cloud controller. We were not able to gain access to the LUKS encrypted device by any means without providing the correct passphrase. All in all, we believe that the approach described here can be safely used to insulate any data uploaded to an encrypted Galaxy instance from malicious access as long as the instance itself and the encryption keys remain uncompromised.

### Cluster support

From the point of view of the IaaS administrator, the option to offer static and/or elastic cluster support to users provides the alternative between guaranteeing a constant pool of resources to those instances attached to a static cluster, or greater control over the efficient usage of the available resources, for those instances that instead rely on an elastic cluster. The latter solution (Fig. 7), in fact, dynamically scales the number of cluster nodes available to a Galaxy instance depending on its workload. From the user point of view, both solutions enable straightforward access to computational resources beyond those assigned to the Galaxy instance virtual hardware, enabling a higher number of simultaneous users, more rapid execution of jobs and additional room for computationally intensive analyses.

**Fig. 7.**
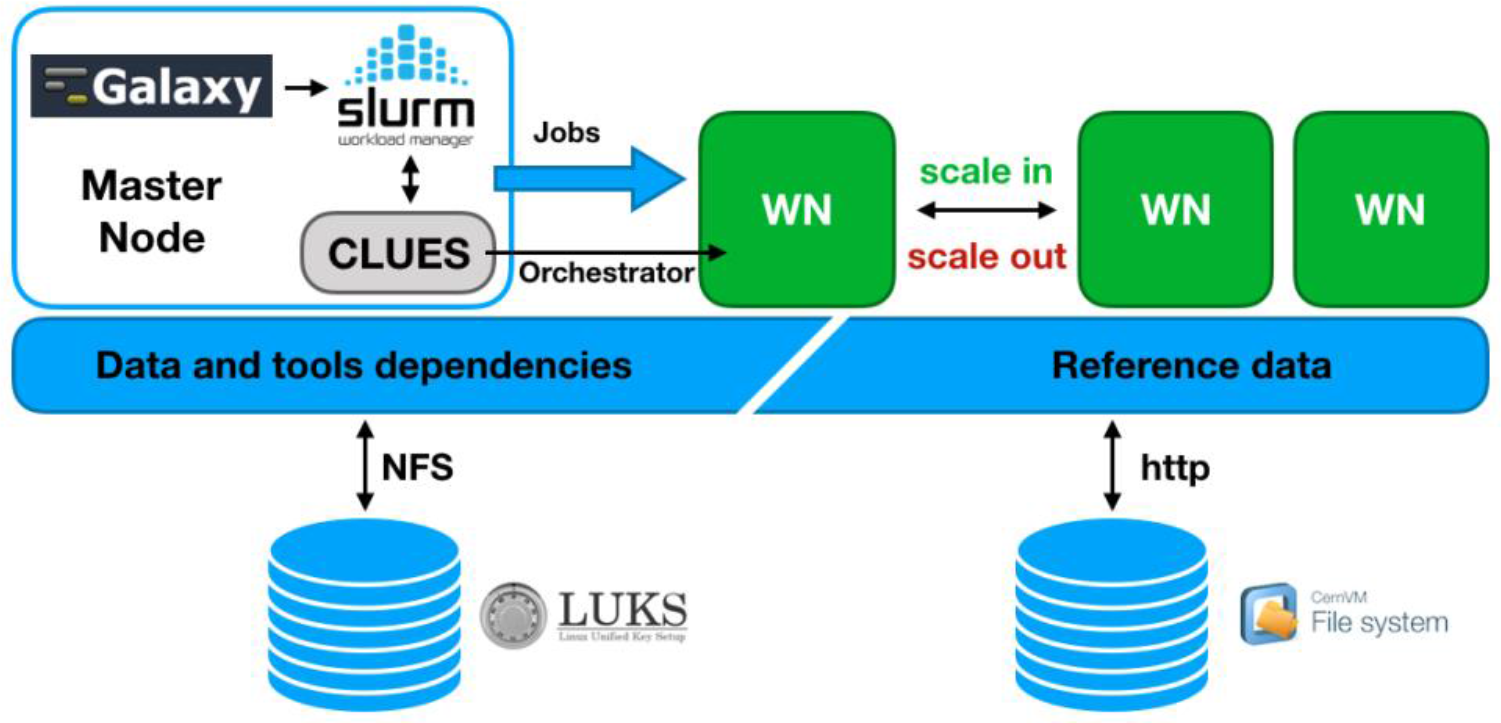
Galaxy elastic cluster architecture. Initially, only the master node, that hosts Galaxy, SLURM, and CLUES, is deployed. The SLURM queue is monitored by CLUES, and new worker nodes are deployed to process pending jobs up to the maximum number set during cluster configuration, thus adapting resources availability to the current workload. The user home directory and persistent storage are shared among master and worker nodes through the Network File System (NFS) enabling the sharing of CONDA tools dependencies. If and when the dependencies are not satisfied by CONDA, the required packages are installed during deployment on each worker node. The CVMFS shared volume is also mounted on each worker node so to ensure that tools have access to reference data.

### Laniakea pilot instance

In order to test and demonstrate Laniakea, we have deployed a pilot service over an OpenStack (Mitaka release) IaaS hosted at the ELIXIR-IT ReCaS-Bari facility [46]. The current INDIGO PaaS layer configuration of this pilot service is reported in Table S4 (Supplementary2.docx). During the first closed beta program, starting Dec 2018, up to 128 CPUs, 256 GB of RAM and 10 TB of disk storage will be reserved for the service. Resources will be gradually increased over the next few months in order to make this prototype instance grow into a full-fledged service provided by the Italian Node of ELIXIR (https://www.elixir-europe.org/). The service front-end is available at elixir-italy-laniakea.cloud.ba.infn.it.

## Discussion

Laniakea offers a feature-rich solution to include a Galaxy on-demand service within the portfolio of public and private cloud providers. This result is achieved leveraging the INDIGO middleware that, having been designed to support a vast array of scientific services, may already be present and supported within the same cloud infrastructure or, as an alternative, can also be used to pilot locally available computational resources from a remote INDIGO deployment. Laniakea’s scalability encompasses a variety of possible Galaxy setups that span from small instances to serve, e.g., small research groups, developers and didactic purposes to production grade instances with (elastic) cluster support enabling multiple concurrent users and computationally demanding analyses. New Galaxy flavours, allowing to implement and make readily available future or existing data analysis pipelines, can be quickly deployed and shared through Ansible recipes to ease the error-prone and lengthy routines of tools installation. We are confident that the Galaxy PaaS delivered with Laniakea can effectively mitigate the need to host and maintain local hardware and software infrastructures in several different scenarios, favouring a more efficient use of the available resources, harnessing the improved reliability offered by cloud environments and also helping to enhance the reproducibility of bioinformatics analyses. Finally, the data security layer of Laniakea addresses relevant issues for the analysis of sensitive data in human genetics research and clinical settings. Future developments will be aimed at improving the compatibility of Laniakea with a broader array of existing cloud setups, and at extending cluster support to other resource managers (e.g., TORQUE [31], HTCondor [47], etc…). Finally, we plan to expand the compatibility of Laniakea with Docker in order to offer Galaxy containers as long-running services, exploiting the great number of dockerized Galaxy flavours already hosted at Docker Hub (https://hub.docker.com/) and maintained by the Galaxy community.

## Conclusions

Laniakea represents a clear example of the current trend of services virtualization, following the direction set forth, e.g., by the European Open Science Initiative (EOSC) Declaration and enables researchers and scientific services providers to implement several recommendations therein outlined. Indeed, Laniakea offers a platform-agnostic, cloud-based service that can be almost effortlessly kept up to date; this, in turn, facilitates the provision of software and services for the Life Science field and beyond. We believe that by easing the barriers posed by the software and hardware layers required to deploy and maintain Galaxy instances, and thus by enabling better access to cutting-edge technology to a broader audience of researchers and other stakeholders, the approach adopted by Laniakea can contribute significantly to the efficient employment of computational resources in the coming years. Finally, Laniakea design could be easily adapted to deploy other tools beyond Galaxy in order to provide additional instruments for the Life Science community or to serve different scientific communities.

## Supporting information

## Acknowledgements

This work has been supported by the European Commission H2020 research and innovation program under grant agreement RIA 653549.

