## Supplementary material for "Laniakea: an open solution to provide Galaxy “on-demand” instances over heterogeneous cloud infrastructures"

| INDIGO Component | URL |
| --- | --- |
| IAM | <https://www.indigo-datacloud.eu/identity-and-access-management>  <https://github.com/indigo-dc/iam> |
| PaaS orchestrator | <https://www.indigo-datacloud.eu/paas-orchestrator>  <https://github.com/indigo-dc/orchestrator> |
| Infrastructure Manager | <https://www.indigo-datacloud.eu/infrastructure-manager>  <https://github.com/indigo-dc/im> |
| CLUES | <https://github.com/indigo-dc/clues-indigo> |
| PaaS Deploy | <https://github.com/indigo-dc/indigopaas-deploy> |
| FutureGateway | <https://www.indigo-datacloud.eu/future-gateways-programmable-scientific-portal>  <https://github.com/indigo-dc/fgAPIServer>  <https://github.com/indigo-dc/APIServerDaemon>  <https://github.com/indigo-dc/LiferayPlugIns> |
| TOSCA repository | <https://github.com/indigo-dc/tosca-types> |
| INDIGO-DataCloud GitHub | <https://github.com/indigo-dc> |
| INDIGO-DataCloud GitBook | <https://legacy.gitbook.com/@indigo-dc> |

**Table S1: URLs of INDIGO-DataCloud software components required by Laniakea.**

| Software | URL |
| --- | --- |
| Documentation | <http://laniakea.readthedocs.io> |
| Encryption script | <https://github.com/galaxycloud-elixir-IT/fast-luks> |
| Indigo-dc.galaxycloud | <https://github.com/indigo-dc/ansible-role-galaxycloud> |
| Indigo-dc.galaxycloud-os | <https://github.com/indigo-dc/ansible-role-galaxycloud-os> |
| Indigo-dc.galaxycloud-fastconfig | <https://github.com/indigo-dc/ansible-role-galaxycloud-fastconfig> |
| Indigo-dc.galaxycloud-tools | <https://github.com/indigo-dc/ansible-role-galaxycloud-tools> |
| Indigo-dc.galaxycloud-tooldeps | <https://github.com/indigo-dc/ansible-role-galaxycloud-tooldeps> |
| Indigo-dc.galaxycloud-refdata | <https://github.com/indigo-dc/ansible-role-galaxycloud-refdata> |
| Galaxy-flavor-recipes | <https://github.com/indigo-dc/Galaxy-flavors-recipes> |
| Reference-data-galaxycloud-repository | <https://github.com/indigo-dc/Reference-data-galaxycloud-repository> |
| Indigo-dc.cvmfs-server | <https://github.com/indigo-dc/ansible-role-cvmfs-server> |
| Indigo-dc.cvmfs-client | <https://github.com/indigo-dc/ansible-role-cvmfs-client> |
| Docker CONTAINERS | <https://hub.docker.com/u/laniakeacloud> |
| TOSCA Template repository | <https://github.com/Laniakea-elixir-it/TOSCA-templates> |
| Instance management script (galaxyctl) | <https://github.com/Laniakea-elixir-it/galaxyctl> |
| PORTAL GITHUB REPOSITORY | <https://github.com/Laniakea-elixir-it/elixir-italy-science-gateway> |
| Running prototype service | [https://elixir-italy-laniakea.cloud.ba.infn.it](https://elixir-italy-science-gateway.cloud.ba.infn.it) |
| VIDEO DEMO | <https://goo.gl/xnWNQd> |

**Table S2: URLs of Laniakea components and documentation.**

| Encryption options | Default configuration |
| --- | --- |
| Cipher | aes-xts-plain64 |
| Key size | 256 bit |
| Hash algorithm | Sha256 |
| Default device | /dev/vdb |
| Device Mapper Name | Randomly generated (e.g. nvdisd) |
| Mount point | /export |
| File system | ext4 |

**Table S3: Storage encryption configuration parameters.**

| INDIGO-DataCloud component | version |
| --- | --- |
| IAM | v1.2.1 |
| Orchestrator | 2.0.0-FINAL |
| Infrastructure Manger | 1.7.6-dev |
| SLAM | v2.0.0 |
| CMDB | v0.4 |
| Monitoring | 1.0.2 |
| FutureGateway Portal | Master branch (customized) |

**Table S4: Current configuration of the Laniakea instance hosted at the ELIXIR-IT ReCaS-Bari facility.**

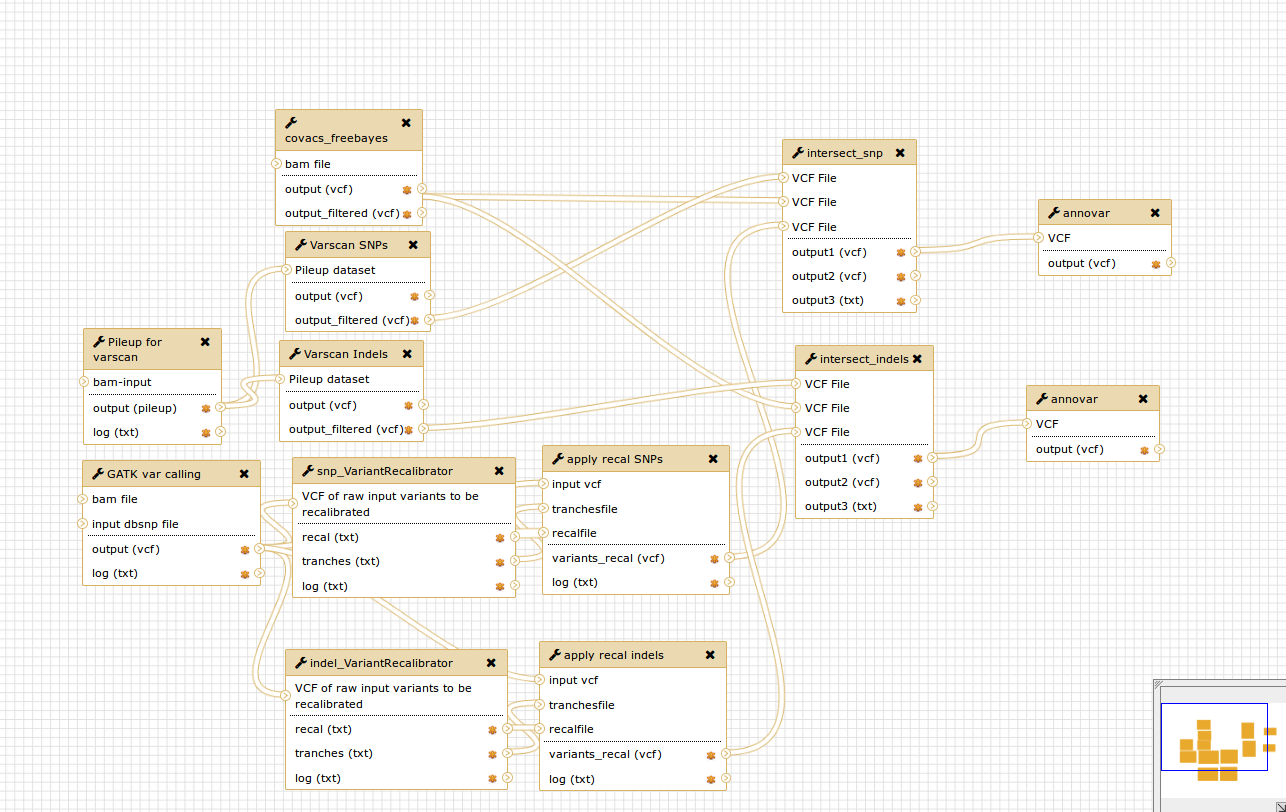

**Figure S1: CoVaCS implemented as a Galaxy workflow for the corresponding flavour.**
