## Supplementary material for "Laniakea: an open solution to provide Galaxy “on-demand” instances over heterogeneous cloud infrastructures"

### **Laniakea data encryption layer**

Device mapper is the Linux kernel driver for volume management and provides transparent encryption of devices through the Linux kernel crypto API, using its device mapper crypt (dm-crypt) module. Dm-crypt is commonly used through Cryptsetup [cryptsetup], a command line interface to dm-crypt, allowing user to setup a new encrypted block device in /dev, specifying the encryption mode, the cipher and the key. Then the device can be formatted with a file system (e.g. ext4), mounted like any other partition and used as persistent storage.

Cryptsetup supports different encryption modes, like plain dm-crypt [cryptsetup] and LUKS volumes [LUKS_web, LUKS_spec] already included in the Linux kernel, but also Loop-AES [loopaes] and TrueCrypt/VeraCrypt [vera] requiring extra modules installation.

We restricted our choice to dm-crypt usage, which exploits Linux kernel built-in APIs, avoiding the installation of any additional external package other than cryptsetup. In particular, the LUKS encryption grants better usability and flexibility to end users without neglecting data security. Unlike others encryption modes, LUKS stores all dm-crypt setup information in the partition header at the beginning of the block device itself, allowing for multiple passphrases that can be changed and/or revoked anytime. It provides robustness against low-entropy passphrases attack using salting and iterated PBKDF2 passphrase hashing.

Cryptsetup allows for different ciphers usage. A cipher consists of three parts: a block cipher, i.e. it is the encryption algorithm, which operate on fixed-length blocks of data; a block cipher mode of operation, which describes how to repeatedly apply a cipher single block operation to data larger than cipher block size and an Initialization Vector (IV) generator, used to randomize the output of the encryption algorithm, ensuring that the same data are encrypted differently with the same key.

LUKS default cipher is aes-xts-plain64, i.e. AES as block cipher, XTS as mode of operation and plain64 as IV generator. The Advanced Encryption Standard (AES) [AES] is a symmetric-key algorithm, I.e. the same key is used either to encrypt and decrypt data, applying several substitution and permutation rounds to plaintext block to produce encrypted blocks. The Xor encrypt xor Tweakable block Cipher (XTS) mode of operation [XTS1, XTS2] is intended specifically to encrypt data on a block-structured storage device, e.g. disk sectors. The mode works with AES as underlying block cipher which is applied two times to each data chunk: the plain text block is combined with the tweak value, i.e. the plain64 IV, encrypted with AES. Then the block is AES encrypted with the key. Finally, the result is combined again with the tweak value before storing the cipher block.

These options represent the current standard on storage encryption and their modification is strongly discouraged, unless user requires particular configurations. For this reason, even if the Laniakea encryption layer can in theory accept user-defined configuration, e.g. different ciphers, we did not expose these options in the user-interface.

**References**

[cryptsetup] <https://gitlab.com/cryptsetup/cryptsetup>

[LUKS_web] <https://gitlab.com/cryptsetup/cryptsetup>

[LUKS_spec] Clemens Fruhwirth, <https://mirrors.edge.kernel.org/pub/linux/utils/cryptsetup/LUKS_docs/on-disk-format.pdf>

[loopaes] http://loop-aes.sourceforge.net/

[vera] www.veracrypt.fr

[AES] Morris J. Dworkin, Elaine B. Barker, James R. Nechvatal, James Foti, Lawrence E. Bassham, E Roback, James F. Dray Jr. Advanced Encryption Standard (AES). November 26, 2001. <https://www.nist.gov/publications/advanced-encryption-standard-aes>

[XTS1] Dworkin, M. NIST Special Publication 800-38E: Recommendation for block cipher modes of operation: The XTS-AES mode of confidentiality on storage devices, Jan. 2010.

[XTS2] Institute of Electrical and Electronics Engineers. IEEE Std. 1619-2007, IEEE standard for cryptographic protection of data on block-oriented storage devices. IEEE

Press, Apr. 2008.
